# Protein Language Model Based Structure-guided Antibody Screening for Disordered Protein Targets

**DOI:** 10.1101/2025.06.21.660895

**Authors:** Akshay Chenna, Prasoon Priyadarshi, Keshav Kolluru, Saurabh Singal, Gaurav Goel

## Abstract

A crucial step in the pathogenesis of Parkinson’s disease involves cell-to-cell transmission of *α*-Synuclein proto-fibrils via endocytosis, driven primarily by the interaction of its disordered C-terminal peptide with domain 1 of Lymphocyte Activation Gene 3 (LAG3) neuronal receptors. High-affinity antibodies have been proposed as therapeutic modalities to delay this progression and subsequent amyloid formation. In our work, we develop an end-to-end computational pipeline to enable rapid screening of antibody sequences that have a high-affinity for the disordered C-terminal peptide of *α*-Synuclein using no information of known binders. This de novo screening was enabled by a structural bioinformatics based *in silico* data generation pipeline combined with a deep learning framework. Our simple feed forward network model built upon sequence embeddings from a protein language model ranked the binding affinities (ΔG) of antibodies to *α*-Synuclein with a high accuracy (Spearman *ρ* = 0.86) when the training and the evaluation datasets contained sequences having some overlap in the complementarity determining regions (CDRs). However, for vastly different CDR sequences, a transformer encoder model trained using the antibody sequence embeddings showed a low Spearman rank correlation of *ρ* = 0.18. The models have a mean Precision@100 of 38 and 12 respectively, significantly outperforming a random process. Overall, our work demonstrates a computational protocol for generating a high quality dataset of antibody-antigen complexes spanning a very large diversity in antibody sequences followed by training of a deep learning model for prediction of high-affinity antibody sequences for a specific protein target with no known binders.

## Introduction

The spread of pathological *α*-synuclein (*α*S) is a crucial event in the progression of Parkinson’s Disease (PD). Recent experimental work has uncovered the mechanism of transmission, propagation and internalisation of *α*S fibrils(1). The interaction of the *α*S fibrils with the neuronal cell surface receptor - lymphocyte activation gene 3 (LAG3), initiates the endocytosis of the assemblies(2) and cell-to-cell propagation. The internalized *α*S act as seeds for auto catalysing aggregation in healthy dopmaine neuronal cells causing cellular degeneration and eventual cell death. The domain 1 of LAG3 (L3D1) is found to preferentially bind the fibrillar forms (K_D_ = 77 nM) over the monomeric states (no binding)(3). Solid state NMR, cryo-EM and biophysical studies revealed strong interactions of the acidic C-terminal of fibrillar *α*S with the alkaline residues of L3D1(4, 5). Inhibition of this interaction is proposed as a therapeutic strategy for Parkinson’s Disease and related synucleinopathies(6, 7). Passive immunization therapies including clinical candidates such as Prasinezumab have shown to inhibit the endocytosis of *α*S, reduce its prion-like spreading and slow the progression of PD(8–10).

In our work, we develop an end-to-end computational protocol to identify antibodies binding to the pathological and disordered C-terminus of proto-fibrillar forms of *α*S to curtail its interaction with L3D1, a key driver of the PD progression. Here, we generate custom datasets of sequentially and conformationally diverse antibodies complexed to the disordered C-terminus of *α*S to train a deep neural network for a direct prediction of the binding energies ΔG from the sequence, with potential applications for rapid *in silico* screening. Our approach computationally mimics the typical development route of therapeutic antibodies that involves discovery *in vivo* via transgenic animals or *in vitro* display platforms, where the initial hits are experimentally matured for affinity enhancement by mutagenesis libraries or via rational engineering of the complementarity determining regions (CDRs)(11). Once high affinity antibodies are identified, they are further engineered to confer drug like properties such as *in vivo* half life and auto-immunity (12).

These conventional experimental pipelines for antibody discovery are time-consuming and resource-intensive and do not provide insights into the epitope/paratope regions and their binding modes. In this work, we develop an *in silico* platform for identification and construction of an optimized library of sequences (pipeline in Figure S1). The procotols developed in this study use deep learning models to perform screening on a sequence library. The deep learning models developed are capable of high-throughput inference in contrast to traditional computational intensive methods such as docking (structure-based). Further, the models developed in this work can be readily extended to guide generative neural networks for the design of paratope regions of the binding antibodies.

## Results

### Selection of structurally diverse and sequentially distinct Fv antibodies

To train an antibody-based neural network model generalizable to the theoretically relevant therapeutic space of antibodies, a large magnitude and diversity of data is assumed(14). To enable this within the feasible limitations of compute requirements, we use a set of non-redundant sequences containing 104K paired-antibody sequences from the Gray Lab(15). This non-redundant set of antibodies was obtained by sequence clustering on the paired sequences from the observed antibody space (OAS)(16). Using the antibody structural predictions from IgFold, we structurally classify each complementarity determining region (CDR) of the antibody using PyIgClassify(17), a database that assigns clusters to CDR conformations using backbone torsions. This results in six cluster assignments per Fv input structure (one assignment per loop). Upon identification of the cluster for the initial set of 104K Fv structures, we selected a minimal set of Fv antibodies that accounted for all observed cluster identites from the inital set as described in the Methods (Section A). This resulted in a set of 8947 Fv antibodies that are both structurally and sequentially as diverse as possible. This minimal set will be used for subsequent data generation pipelines.

### Generation of alternate conformations of the Fv antibodies

The structures of the Fv domains of the antibodies predicted using protein structure prediction models are independent of the context of the antigen. The CDRs of the antibodies are often flexible and can undergo significant conformational changes upon binding to the antigen(18, 19). A study by Guest et al.(20) comprehensively characterized and compared the structural changes of the antibody paratopes and the CDR loops between the *apo* and the *holo* states. Significant conformational changes in the interface (RMSD > 3.5 Å) and loop backbones (RMSD > 6 Å) were observed. Since protien structure models are inadequate at predicting alternate conformations of CDR loops and inter chain (V_H_-V_L_) orientations of the antibody(21, 22), we have used explicit-solvent all-atom molecular dynamics simulations to capture alternate but functionally relevant states of the antibody. In order to expedite the sampling of conformations, we relied on simulations at temperatures elevated from the physiological conditions. To evaluate the sampling temperatures that balances structural diversity with physicality, we performed 0.5 µs long simulations of G6 Fv (binds to VEGF) starting with the *apo* state (PDBID: 2FJF) at temperatures of 310 K, 340 K, 370 K, 450 K as detailed in the Methods Section B. We compare the generated ensembles to the conformations derived from a Markov-state model (MSM)(13). We notice that (Figure 1A) the trajectory simulated at the highest temperature (450 K) results in CDR L2 conformation that was closest to all the MSM-derived microstates. CDR L2 was chosen for preiliminary comparison as the G6 Fv is known to undergo a large conformational change > 7 Å during the *apo* to *holo* transition. This *holo* conformation was found by Fernandez-Quintero et al. to be sampled in the *apo* trajectory over the course of 18.6 µs MD simulation. Similar results are observed for the remaining CDR loops (Figure S2). Further, we evaluated if sampling at higher temperatures for a shorter duration (here 50 ns) yield conformations different from those obtained at physiological temperature (310 K). We simulated 16 different Fv antibodies *apo* taken from Guest et al.(20) and compared the ensemble similarity of trajectories generated at higher temperatures to the baseline simulation (500 ns, 310 K see Methods). On comparison with this baseline, we observe that the trajectory simulated at 450 K yields an ensemble that is most diverse (Figure S3). In addition, we observed that a further shorter trajectory generated for 10 ns when clustered using a cutoff of 1.5 Å yielded an ensemble that is closer the MSM microstates than a longer trajectory (here 50 ns) clustered at 2.5 Å (Figure 1B). Thus we concluded on the use of 450 K for sampling alternate conformations followed by clustering the trajectory using a backbone cutoff of 1.5 Å on the CDR residues. The conformations of the Fv region of an antibody clustered from a molecular dynamics trajectory are depicted in Figure 1C.

**Fig. 1.**
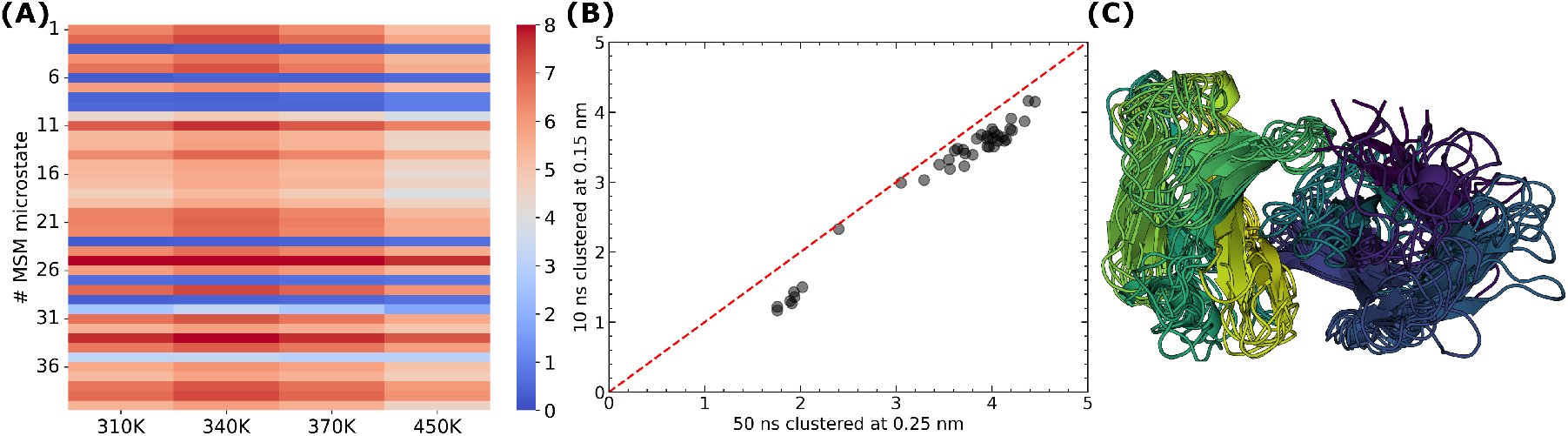
Sampling of alternate backbone conformations and comparison of CDR backbone RMSDs of short *apo* trajectories simulated at different temperatures to the microstates derived for G6 Fab from Fernández-Quintero et al.(13) **A** The heatmap depicts the minimum backbone RMSD of CDR L2 of *apo* G6 (binds to VEGF) simulated at different temperatures for 50 ns to microstates derived from a Markov-state model (MSM) analysis on a 18.9 µs long aggregate simulation. Plots for the remaining CDRs are shown in Figure S2. **B** The CDR L2 RMSDs of the cluster centers derived from clustering a 50 ns trajectory using a cutoff of 2.5 Å on the backbone with respect to the MSM microstates are compared to a 10 ns trajectory but using a stringent cutoff of 1.5 Å. Both trajectories were simulated at 450 K. **C** A solution state ensemble of the *apo* Fv antibody clustered from a 10 ns trajectory simulated at 450 K used for docking to *α*-Synuclein (*α*S) and subsequent affinity maturation.

### Resolving the bound conformation of the disordered C-terminus of *α*S

*α*S is a disordered protein in its monomeric state. *α*S oligomerizes to form proto-fibrillar or fibrillar assemblies that are pathogenic as they are known to aid in inducing oligomerisation by secondary nucleation (23). *α*S proto-fibrils are known to infect healthy neurons by interacting with a neuronal cell surface receptor called LAG3(1). However, an experimentally resolved structure of *α*S complexed with LAG3 is not determined. Here, we use the NMR chemical shift deviation and 2D HSQC data from Zhang et al.(4, 5) to model the bound conformation of C-terminal peptide *α*S (residues 118-140) to the domain 1 of LAG3 (L3D1). This disordered peptide was docked to L3D1 using these NMR data as restraints on the possible receptor-peptide contacts (Methods Section C). The structure of L3D1 was obtained from the predictions of AlphaFold (Figure S5), while the conformations of *α*S were taken from the proto-fibrillar assembly(2N0A)(24). Figure 2 depicts the fraction of contacts made by both the *α*S peptide and the L3D1 receptor in the docked poses when compared to the NMR derived contact information. Each point represents a unique docked pose whose size determines the docking score (larger sizes correspond to better docking scores, here HADDOCK) and the darker shades correspond to a higher number of docked poses with the respective contact fraction. While, during each docking run 50% of the input restraints are randomly removed, a majority of the docked poses converge to a near-native ensemble of structures. The docking results also indicate that the complexes displaying a higher fraction of NMR contacts (fraction > 0.5) are more stable (from HADDOCK scores). Six structurally distinct conformations of *α*S (Methods Section C) were obtained by clustering (Figure 3A,B) and were used as the target set of antigens.

**Fig. 2.**
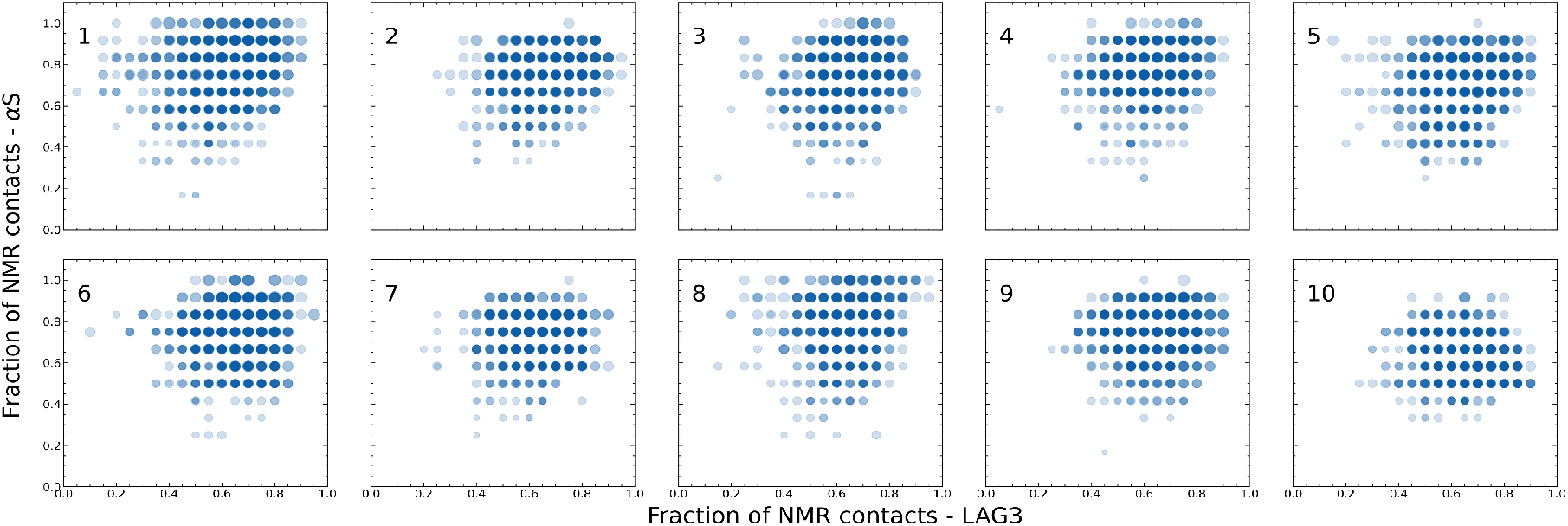
Contact similarity of the docked LAG3-*α*S complexes to NMR determined contacts. Docking simulations were performed for LAG3 Domain 1 and the C-terminal residues of *α*S using the NMR derived contacts as restraints from Zhang et. al. (4, 5). The fraction of contacts common between NMR and the docked poses is plotted for LAG3 and *α*S. Each point represents a docked pose whose size is indicative of the docking score, larger sizes implying better HADDOCK scores. Darker shades indicate more populated states.

**Fig. 3.**
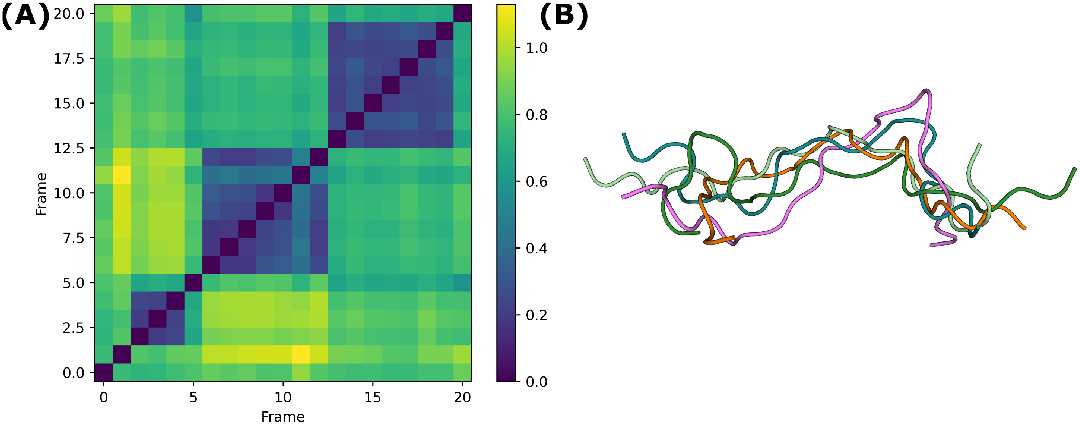
Ensembles of *α*S and Fv region of the antibody. **A**. A matrix of the backbone RMSDs (in nm) of the C-terminal peptides (residues 118-140) of *α*S with the highest NMR determined contact similarity to LAG3 and binding affinities. **B** A visual depiction of the starting ensembles of *α*S obtained after backbone clustering.

### Generation of synthetic training data of Fv-*α*S bound poses

To enable structure based screening, we created a synthetic dataset of Fv antibodies bound to either of the six conformations of *α*S. For each Fv antibody sequence (8947 sequences), we generated an ensemble of alternate conformations using the molecular dynamics simulation protocol and CDR backbone clustering discussed previously. We generate an ensemble of 67K conformations starting from 8947 *apo* structures. Figure 1C depicts an ensemble of conformations for a particular Fv antibody after MD simulations and clustering. This ensemble of Fv antibodies was used to generate an ensemble of Fv-*α*S docked complexes. Since most of these complexes are not expected to bind *in vitro*, we use the clustered docked complexes (see Methods Section D) to redesign the paratope to exhibit enhanced binding affinity to *α*S. We use RosettaAntibodyDesign (RAbD) to perform *in-silico* affinity maturation. The docked poses are taken as the input and mutations on the CDRs in contact with the antigen (*α*S) yielding sequentially diverse structuers with improved affinities, mimicking *in vivo* somatic hypermuations. Figure 4A highlights the improvements in the binding affinities starting from the poses dervied from docking (in *⋆*). Most starting docked poses show an improvement in both the absolute binding affinities (ΔG) as well as an overall improved interface (ΔG per buried interface area). These complexes are expected to exhibit a tighter fit with the disordered *α*S(Figure 4B) and thus micmic an experimental dataset derived from yeast display technique. Figure 4B additionally highlights the conformational sampling of the CDRs and the docked *α*S peptide upon *in silico* affinity maturation. Using RAbD, we have generated over 250,000 complexes whose Fv sequences and the corresponding binding affinities (in Rosetta Energy Units (REU)) will be used to train a sequence-to-affinity model

**Fig. 4.**
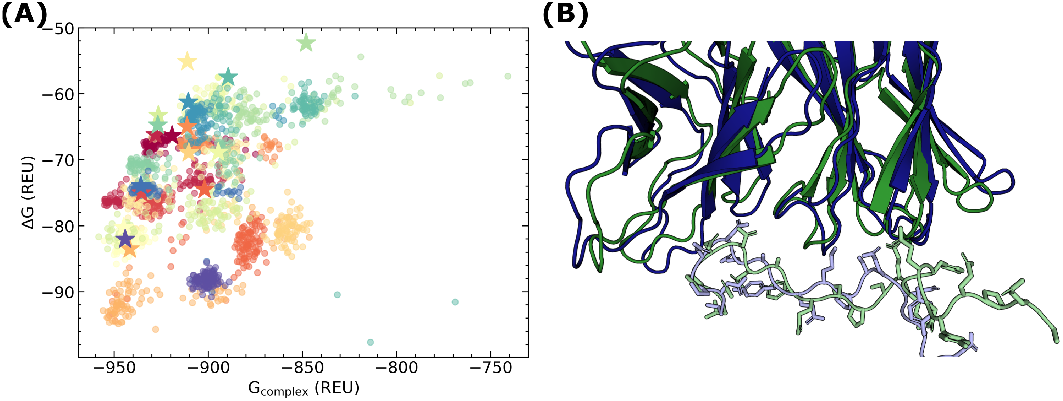
Paratope redesign of the Fv antibodies in the *holo* state. **A** The docked poses of the Fv antibody and *α*S obtained after clustering using a fraction of common contacts (FCC) criteria of 0.6 are used as an input to RosettaAntibodyDesign (RAbD) for improving the binding energy by modification of the paratope residues. The rosetta energies of the complex (G_complex_) plotted against the binding energies (ΔG) are shown. The starting docked structures are depicted with a *⋆* and the complexes derived from each docked pose are colored accordingly. **B** A depiction of the conformational changes of the CDRs upon paratope redesign. The docked complex is colored in green and the affinity matured complex is colored in blue. The antigen (*α*S) is shown in lighter shades with side chains.

### Evaluating model performance at predicting the binding affinities from Fv sequence

Using the dataset of the antibody-*α*S complexes with the corresponding binding affinities, we aimed to train a neural network with the capability of accurately predicting the binding affinities of the antibodies with *α*S using only the Fv sequence as the input. We use deep-learning based protein lanugage models as the basis for making sequence-to-affinity predictions. The neural networks were trained and evaluated for predicting the ΔG of Fv-*α*S binding. We evaluated the predictions from a simple feed forward network and a ESM-2 based finetuned model5A,B. Using the sequence averaged embeddings from Evolutionary Scale Modelling (ESM-2) with original weights(25) and a model fine-tuned on the paired input Fv sequences a feed forward neural network was trained to predict the ΔG (see Methods E). We observed high Spearman rank correlations (*ρ*, Figure 5C and Figure S6) of 0.79 and 0.86 respectively (*ESM2 MLP* and *ESM2 FT*) on the test set consisting of about 25K Fv sequences. Additionally, we found the model to make highly precise predictions, correctly identifying 38 out the best known 100 Fv sequences from its top 100 predictions (Precision@100 = 0.38), while the probability of obtaining at least 38 correct points by chance is extremely low (10^−67^) 5C. Importantly, we note that our model is capable of identifying high-affinity sequences from a set containing similar CDR sequences (>70% sequence identity) (Figure 5D). Figure 5D plots the distribution of the scores of the top 100 sequences predicted by the *ESM-2 FT* model in blue, while the distribution in orange are the sister sequences of those top 100 sequences, i.e. those derived from the same parent upon RAbD maturation.

**Fig. 5.**
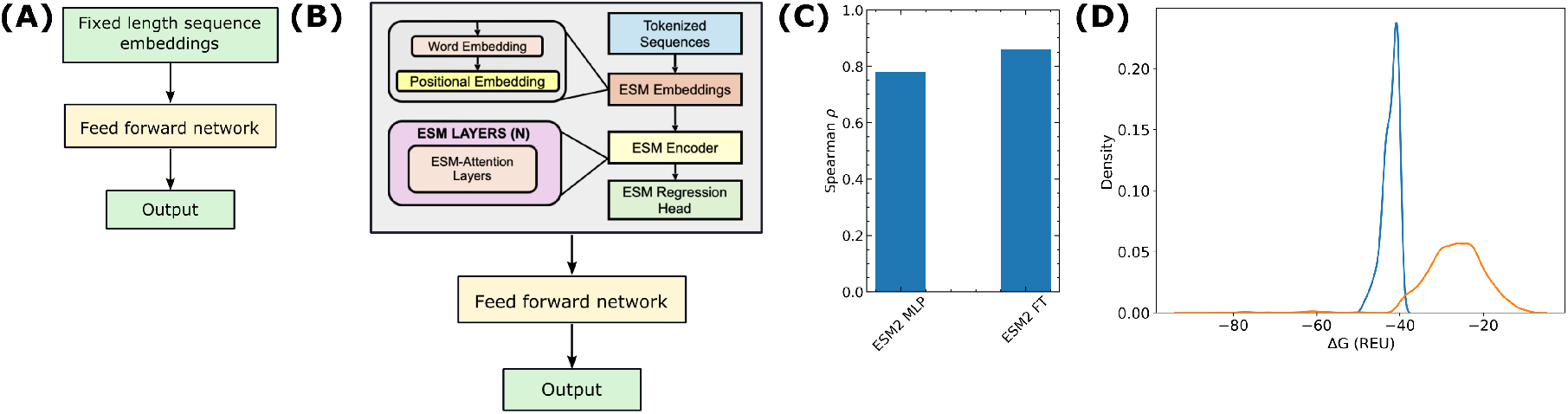
Model architectures and evaluation of the model predictions for the ΔG of Fv-*α*S binding. **A** Fixed-length embeddings from the ESM-2 (150 M) language model are used as an input to a feed forward network. This model is abbreviated with *ESM2 MLP*.**B** For training the *ESM2 FT* model, the ESM-2 (150M) architecture is fine-tuned with the sequences of antibodies generated via the RAbD pipeline to generate contextual embeddings. **C** The spearman correlation coefficients (*ρ*) for the binding affinity predictions on the test sets are shown for the *ESM2 MLP* and the *ESM2 FT* models. **D** The distribution (in blue) of the best 100 ΔG predictions from the *ESM2 FT* model compared against all the scores of all their sister sequences in the training dataset (in orange).

Secondly, as a rigourous test of our model in its ability to predict the affinity of *de novo* sequences, we evaluated the prediction of ΔG on a train-test split sharing sequences with < 20% CDR sequence identity. We evaluated several architectures and protein langugage models for this task (Figure 6A and Figure S6). For this case, where the test and the training set CDR sequences share low sequence identities, we find that the MLP model using vanilla the ESM-2 fixed length embeddings fails to rank the sequences correctly (Figure 6B). However, the ESM-2 model fine tuned to our dataset (ESM2 FT) of paired sequences gives a slight improvement of Spearman *ρ* = 0.13. We further explored if the ESM-2 model fine-tuned to all the paired sequences from the Observed Antibody Space (ESM2-Paired MLP) performs better. ESM2-Paired MLP was found to be equivalent (Spearman *ρ* = 0.12) to the *ESM FT* model, indicating no additional benefits of training on the entire paired OAS sequences. In addition to the embeddings from ESM-2, two models that have encoded structural features (AntiBERTa2 and ProtT5-XL) have been evaluted, both models give a low-moderate predictions of Spearman *ρ* of 0.1 and 0.15 respectively. Finally, we used the per token embeddings of the CDR sequences as an input to two transformer encoder layers with multi-head attention (Methods Section E), to predict the binding affinities (model architecture: Figure 6A). The Sperman *ρ* was observed to be the highest from all the models we have tested. We also observed that 12 out of the top 100 model predictions coincide with the best 100 known Fv sequences in the test set, indicating a precision of 0.12 in the top 100 results (random probability = 10^−15^) (Figure 6C).

**Fig. 6.**
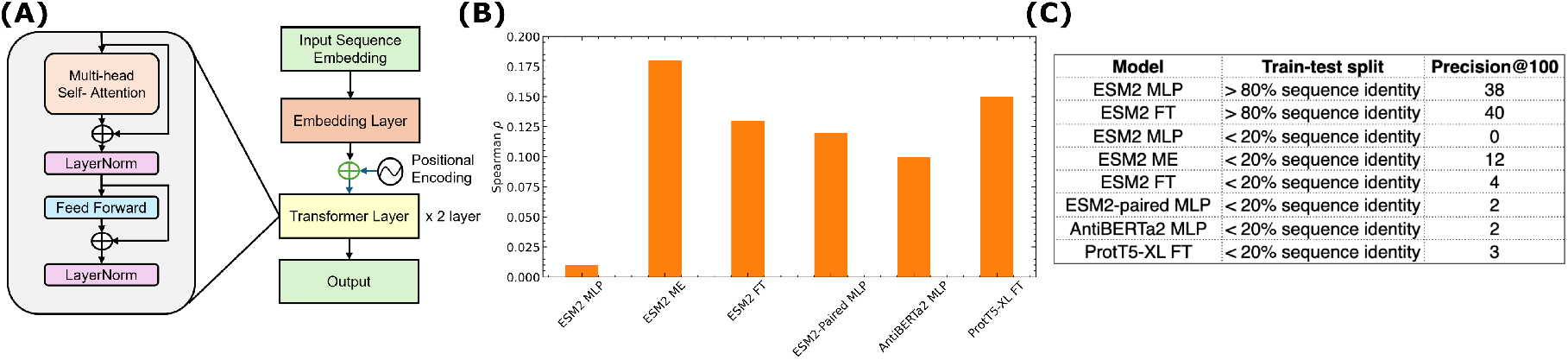
Model architectures and evaluation of the predictions of ΔG of Fv-*α*S binding for out-of-training-distribution antibody sequence (<20% CDR identity) **A** The neural network architecture for the the Model Extension (*ESM2 ME*) network is depicted. The sequence embeddings generated per-residue are re-learnt by the transformer encoder architecture before generating a fixed-length embedding to predict the binding energies (ΔG). **B** For the training-test split with low CDR sequence identity (<20%), the spearman *ρ* is compared for different model embeddings and architectures. **C** The model-wise precision of the top 100 predicted sequences compared against different train-test split ratios based on the % CDR sequence identity evaluated for all the models depicted in Figure 5C and 6B.

## Discussion

In our present work, we developed an end-to-end computational pipeline for training a deep-neural network capable of predicting the binding affinity to a disordered protein target (the C-terminal peptide of *α*-synuclein) from the Fv sequence of the antibody alone using the binding information of the antibody-antigen complex as the training set. We showcase a detailed and generalizable approach for the generation of a synthetic dataset of the complexes of the Fv-antibody and the protein target *in silico* for training a neural network based on a protein language model (PLM). We find that the model generalizes and ranks the sequences accurately when the sequences in the test set do not vary significantly (>70% CDR sequence identity) from the training set. The performance of the model significantly drops when the datapoints in the test-train split differ significantly (< 20% CDR sequence identity), while displaying orders of magnitude higher accuracy than a randomly sampled set.

Here, we present an *in silico* data generation pipeline that may serve as an alternative to intensive experimental efforts that often rely on deep mutational scans or library based assays(26). *In silico* data generation approaches offer several benefits that are not accessible by wet-lab assays. For example, a computational approach enables a high-throughput data generation with detailed resolution of the antibody-antigen complexes and the binding interface(27). In particular, our approach allows us to explicitly model alternate conformations of the antibody that have been largely ignored in most recent structure-based studies. Using solvent molecular dynamics we were able to generate a diverse ensemble of CDR loop conformations and heavy-light chain orientations (Figure 4C) that can allow improved modelling of the downstream processes such as docking and paratope re-design (Figure S7). We find that a short MD trajectory at elevated temperatures generates significant diversity in the conformations and these conformations are found to more faithfully capture the microsecond dynamics of the loop-motions of the Fv CDRs. By relying on multiple conformations instead of a single state predicted to be the ground truth by protein folding networks, we alleviate the risk of relying on low-confidence predictions. Additionally, we find that generation of alternate states using MD results in a more diverse ensemble than methods relying on subsampling AlphaFold2(28, 29) or its extensions via enhanced sampling (30) (data not shown).

Our approach is well-suited for disordered protein targets as we inherently model protein flexibility during docking and design. Despite the structure of the C-terminus of *α*S bound to its cell surface receptor (domain 1 of LAG3) not being experimentally determined, we used information of the contacts made at the interface of the complex to guide the docking process and computationally resolve the bound state conformations of the disordered *α*S peptide. We find that docked poses with a higher fraction of native contacts are energetically more favourable (2). From the docked complexes, we find multiple yet diverse conformations of *α*S that mimic the NMR derived native contacts with high fidelity, highlighting the disordered and fuzzy nature of *α*S being maintained in the complexed state. Multiple top ranked diverse conformations were used as target antigens for docking with random Fv antibodies despite no evidence for the epitope recognition by the paratopes. To address this non-nativeness challenge we optimized for the binding by re-designing the paratope residues on the Fv antibody. This process of *in silico* maturing the affinity of the antibody is reminiscent of somatic hypermutations on the germline antibody sequences. The sequence re-design steps also address the limitation of the lack of sampling space explored during docking as highly flexible proteins (here, both the antigen and the antibody CDRs) exhibit larger degrees of both the rigid body and conformationaly degress of freedom(31). The sequences obtained post *in silico* affinity maturation involve mutations at multiple sites on the paratope concurrently. Such a diverse exploration of the sequence landscape is not achieved by experimental techniques that are often limited to single-point mutations(32, 33). Our approach accounts for an important feature of the datasets used for training as higher-order mutational effects are known to alter the fitness of the sequence landscapes of the antibody-antigen binding via epistatis (34). These complex landscapes when not represented in the training set pose a significant challenge to downstream model training for prediction tasks (35).

Here, in our work, for the first time, we demonstrate a deep learning based approach designed to rapidly predict the ΔG of antibody binding to disordered targets, where the antigen natively forms a fuzzy complex with its receptor (L3D1). Using generalized and antibody specific protein-language models as the basis, we train shallow and deep neural networks for learing the sequence to affinity relationships. Our results show that while accurate predictions are possible for similar CDR sequences (>70% sequence identity), the model struggles to make accruate predictions for cases with a large deviation (<20% sequence identity). This demonstrates the ability of our model to accurately predict the binding affinities of naturally somatically hypermutated antibodies to aid in the discovery of therapeutic antibodies, where the CDR sequence identities with respect to the germline are usually at least 70%. In addition, our model can be used as an orcale for scoring the outputs of design algorithms for rapid high-throughput optimization around a starting sequence. The inability of our model to predict the affinities of *de novo* sequences is in line with the recent work by Greenshields-Watson et al., where the authors find that deep learning models are unable to make out-of-domain predictions of CDR properties and that even minimal amounts of data of closely related CDRs can significantly improve predictions (36). Similarly, the importance of the volume of data and the diversity in the training dataset has been shown to improve a generalizable model for ΔΔG predictions (14). Most recent works that rely on predicting the binding affinities of proteins and antibodies have shown to make reliable predictions of the effects of mutations rather than for *de novo* sequences (37, 38).

Our results indicate that a generalizable model to predict the properties of antibodies (here ΔG) can only make reliable predictions when the training set contains diverse sequences and structure beyond the ones we have chosen. This is understood by the fact that the over 10^12^ unique and exceptionally diverse antibodies are present in the human repertorie(39) which further undergo variations based on CDR hypermutations. Only a small fraction of these are captured in existing databases(40). Neural network predictions relying solely on language models for antibody property predictions are inferior as compared to other classes of proteins as the sequence relationships of the CDRs are not conserved by evolution (25, 38). However, as demonstrated by experimental efforts using Trastazumab as a test system, Chinery et al.(41) and Bachas et al.(42) clearly show that pipelines that parallel our computational approach can be useful in improving already known antibody binders by improving classification accuracy or by sequence design. Similarly, as demonstrated, the utility of our model lies in making incremental mutations yet yielding designs with vastly superior functions not found in nature(43, 44). Our model can also serve as a foundation for improving fitness via Fv sequence evolution as demonstrated by Hie et al.(**?**). Some pertinent future directions will involve further improvement of the model by fine-tuning and incorporating the high resolution structural information of the antibody and antigen complex (45, 46) or using sequence-to-structure tokens as inputs to the neural network(47, 48). Other ways of fine-tuning or re-training the base language models include using re-inforcement like approaches such as Direct Preference Optimization (DPO) that do not overfit the model to the training data(49) or using the biophysical information from the simulations(50). In fact, in the recent work of Widatalla et. al. (49) the ESM language model fine-tuned using (DPO) a Spearman *ρ* of about 0.2 was found on the AB-Bind dataset(51), consisting of the binding affinities of antibody mutants with the antigens thereby indicating the potential saturation in the accuracy in ranking mutations of current state of the art neural network models.

## Methods

### A. Selection of sequentially and structurally diverse variable (Fv) regions from the observed antibody space (OAS) (16)

A set of 104 K non-redundant paired antibody sequences, clustered using MMseq2(52) using an identity cutoff of 0.95 on the paried antibody sequences from OAS were taken from the predictions of IgFold(15). To evaluate structurally diversity of the starting CDR loops conformations of the heavy and light chains, we used the Rosetta implementation of PyIgClassify (17, 53). For each Fv antibody, a total of six cluster assignments are output (one assignment per CDR) by PyIgClassify. Upon cluster identification, we selected the minimum number of Fv antibodies that accounted for all the cluster assignments from the initial antibody dataset (104K structures).

### B. Generation of an ensemble of solution-state conformations via explicit solvent MD at elevated temperatures

To generate alternate conformations sampling the CDR loops and the V_H_-V_L_ orientations, we performed molecular dynamics simulations at multiple temperatures. For a set of 16 antibodies with both the *apo* and *holo* conformations resolved(20), we performed 500 ns long simulations starting from the *apo* conformations (from the PDB) at temperatures of 310 K, 340 K, 370 K, 450 K in explicit solvent. Using the simulations at 310 K as reference, we calculated the diversity in the loop conformations generated at the elevated temperatures (340 K, 370 K, 450 K). The diversity was evaluted by measuring the fraction of trajectory frames simulated at physiological temperature that are structurally similar to those at elevated temperatures. CDR backbone RMSD of less than 2 Å was used as a criteria for structurally similarity. Further, for the 2FJF Fv antibody, shorter MD simulations (at all four temperature levels) of 50 ns were used to compare the CDR loop structural similarity to the microstates derived from a Markov-state model (MSM)(13). Finally, we compared the CDR loop structural similarity for an even shorter trajectory of 10 ns clustered using 1.5 Å backbone RMSD (*gromos* clustering algorithm) to the MSM microstates. Having found that a 10 ns simulation at an elevated temperature of 450 K generated structures closer to the MSM microstates (than a 50 ns trajectory at 450 K, clustered at 2.5 Å backbone RMSD), the diverse antibodies selected in the initial stage were simulated and clustered as per these conditions to obtain an ensemble of conformations for docking. All-atom molecular dynamics (MD) simulations using the CHARMM-36m forcefield (54) were performed for the *apo* Fv antibodies poses generated by IgFold and filtered in the previous step. Na^+^ or Cl^−^ was added to maintain charge neutrality and the system was dissolved in an aqueous solution of NaCl 0.15 m. Following a series of equilibration steps, a 10 ns MD simulation was run under NPT conditions, with coupling to a Nose-Hoover thermostat (*τ*_*T*_ = 4 ps) and a Parrinello-Rahman barostat (*τ*_*P*_ = 5 ps). This trajectory was corrected for periodic boundary conditions (to fix broken molecules and keep the heavy and light chains intact). The parameters and scripts for running molecular dynamics trajectories and clustering are available in the supplementary material.

### C. Docking of C-terminus peptide of *α*S with the Domain 1 of LAG3 (L3D1) using experimentally derived contact restraints to obtain *α*S conformations as natively bound to L3D1

The initial structure of the Domain 1 of LAG3 (L3D1) protein was obtained from the predictions of AlphaFold2 implemented via ColabFold(55). The sequences of residues 23 to 167 of murine L3D1 and the C-terminus peptide *α*S of (residues 118-140) were input together to ColabFold(56). The structure of L3D1 with the highest pLDDT (and low pAE, Figure S4) was chosen as the starting pose for docking. For *α*S, the 10 conformations of the the C-terminal residues (118-140) were taken from proto-fibrillar assembly (PDB:2N0A)(24).

Information-driven docking was performed using HADDOCK to obtain the docked poses(57). The L3D1 residues with high 1H-NMR chemical shift deviations upon *α*S binding (R58, G59, G60, V61, I62, W63, A101, G103, L105, R106, S107, T146, R148, R152, A153, L154, S155, C156, S157 and L158) were chosen as input restraints on L3D1(5). While residues D121, N122, E126, S129, E130, G132, Y133, Y136, and A140 on *α*S were used as contact restraints, as determined from 2D HSQC spectra(4). 5000 rigidly docked poses were generated starting from each *α*S conformation with subsequent refinement in explicit water using the recommended HADDOCK parameters. *α*S conformations from the docked poses with the fraction of NMR derived contacts > 0.5 (both for *α*S and L3D1) were sorted as per the HADDOCK scores and the top 10% of poses with the highest boltzmann weights were selected for clustering. The conformations of *α*S were clustered using its backbone atoms and a cutoff of 1.5 nm was applied.

### D. Generation of Fv-*α*S docked complexes and Fv paratope re-design via affinity maturation

The conformations of *α*S obtained upon clustering (from the *α*S-L3D1 complex) were used to dock against the ensemble of antibodies (obtained from short MD simulations at elevated temperatures, followed by clustering the CDRs). For *α*S, the residues used as restraints for docking to L3D1 were re-used here. While for the Fv antibodies, the docking was restrained to only the CDR regions. For identification of the CDR regions, the Fv antibodies were renumbered using the Chothia scheme(58) using the ANARCI tool(59). Following the renumbering PyRosetta was used to identify the CDR loops. These residues were used to generate the restraint files for docking. HADDOCK was used to generate the docked poses. Rigid body docking was performed and 100 docked poses were generated for each conformation of the antibody. For a given Fv sequence, a conformation of *α*S was randomly chosen from the earlier clustered conformation set for docking. The recommended parameters were used for executing HADDOCK. The antibody poses were clustered using the fraction of common contacts (FCC) of 0.6(60). This clustering was performed for each Fv sequence, i.e. by considering all docked poses generated from a given Fv sequence.

The cluster centers were renumbered from the Chothia scheme to AHO scheme prior to Fv paratope redesign using Rosetta’s numbering scheme converter. The Fv-*α*S complexes were relaxed using the RosettaRelax application(61). Following RosettaRelax, the light chains of the antibodies were identified using ANARCI before inputting to RosettaAntibodyDesign (RAbD)(62). RAbD was used to generate mature sequences of the antibody paratope that confer improved binding affinity to the epitopes on *α*S. A sampling temperature of 7.5 k_B_T was used. 25 cycles for the outer loop responsible for randomly choosing a CDR loop were applied. One inner loop was run for each outer loop, where the sequence and the structure of the CDRs were optimized for binding affinity. The RAbd protocol is detailed in the supplementary material. Starting from the docked poses, a total of 250K complexes were generated.

### E. Training neural networks for prediction of binding affinities

#### Training using fixed-length embeddings

Embeddings from Evolutionary Scale Modelling (ESM-2) (25) were used for training a neural network to predict the binding affinities (ΔG) from the sequence. The general-purpose embeddings on the entire Fv region (heavy and light chains paired together) were taken from the final layer of the 48 transfomer model trained with 15 billion parameters. Per-residue embeddings (a 5120 length vector) were generated for each of the RAbD-designed sequences. For a given sequence, the per-residue embeddings were averaged over all residues (mean pooling) to yield a fixed-length embedding with a dimensionality of 5120 features. The fixed-length per-sequence embedding was input to a feedforward neural network with two hidden layers of 512 nodes, displaying the binding energy (ΔG) in the last layer (1 node). A GELU activation function was used in between the connections to introduce nonlinearity. We used a train-test split of 0.9:0.1. The mean squared error of the predicted binding energies was the loss function. Regularization was introduced by dropping 20% of the nodes in each layer. All data were standardized and normalized after each layer. Training was performed for 10,000 epochs (using a single batch). The weights were optimised using the Adam’s optimizer with a learning rate of 0.0001. To evaluate our predictions, we assess the overall goodness of fit to the truth of the ground by the coefficient of determination for ΔG of the given sequence (*R*^2^) and the Spearman correlation coefficient for the rank (rank 1: most negative ΔG) of the given sequence (*ρ*) (see figure 5A). A similar protocol was followed using the embeddings from an ESM-2 model fine-tuned on Paired antibody sequences(63) and a structure-informed language model AntiBERTA-2CSSP(21). These models are called suffixed with “MLP".

#### Model Extension of ESM-2 with a transformer network (ESM-2 ME)ss

To generalize the training to better capture the sequence diversity, a transfomer-based architecture was explored to account for inter-residues dependencies. The ESM-2 model with 640 embedding dimensions per token (output from 30 transformer encoder layer model) was used as input. The positions of the residues obey the Chothia numbering scheme. The input sequence was positionally encoded before inputting it into the encoder layer. The encoder layer has a hidden size of 256 with two attention heads. The encoded output was pooled using the attention weights learnt from the transformer layer. This conversion to a fixed-length embedding was then forwarded to a two-layer fully connected neural network with 64 nodes each. A differential spearman rank correlation coefficient was used as a loss function (64), and the weights were optimized using the Adam optimizer. The architecture is highlighted in figure 5B (ESM2 ME).

#### Fine tuning protein language models

The following architectures were used for fine-tuning existing language models for prediction of the binding affinities:

##### ESM-2 fine tuning (ESM2 FT)

The fine-tuning process is configured using the ‘TrainingArguments’ class from the Hugging Face Transformers library. The training arguments include the number of training epochs (set to 60), batch sizes for training and evaluation (set to 8), warmup steps (set to 500) to ensure learning rate will be gradually increased over the first 500 training steps for a stable and smooth start to the training process., weight decay (set to 0.01) to avoid overfitting, logging settings, and evaluation strategy (set to “epoch”). The model architecture is highlighted in figure 5C (ESM2 FT)

##### ProtT5-XL fine tuning (ProtT5-XL FT)

Training metrics are logged every 100 steps, providing regular insights into the model’s performance. The learning rate is set to 3e-4 (0.0003), which is a standard value for ensuring steady convergence. The ‘AdamW’ optimizer, which is the default in Hugging Face’s Transformers library, is used for optimization. To manage the batch size within hardware constraints, gradient accumulation is employed; the batch size per device train is set to 8, with gradient accumulation steps of 4, effectively making the batch size 32. The model is trained for a total of 5 epochs, balancing between sufficient training and risks of overfitting(65).

## ACKNOWLEDGMENTS

We thank KnowDis Data Science LLP for providing partial financial support for conducting this research. We thank FITT, IIT Delhi for providing project management support. We thank IIT Delhi HPC facility for computational resources. AC thanks Prime Minister’s Research Fellowship for the scholarship.

