## Supplementary Text for "Protein Language Model Based Structure-guided Antibody Screening for Disordered Protein Targets"

### Supplementary Information for

#### Development of an *ab-inito* protocol for structure-based antibody screening targeting intercellular transmission of proto-fibrillar $\alpha$-Synuclein

**Akshay Chenna, Prasoon Priyadarshi, and Gaurav Goel**

**Gaurav Goel**
****

**This PDF file includes:**

Supplementary text
Figs. S1 to S7
SI References

#### Supporting Information Text

The following sections contain details relevant to the protocols developed in this study and the data generated from simulations and neural network models.

#### 1. Code Execution, packages and dependecies

**Code Execution.** Relevant code is available on Github. To reproduce the results, the sequence of scripts is:

- Identify CDR clusters: run_cluster_identify.sh
- Select distinct clusters: cdr_analysis.ipynb
- MD of antibodies: md_production.sh (with md.mdp), correction_md.sh.
- Cluster MD poses: cluster_md_apo.sh
- Docking restraints: find_cdrs.sh (with find_cdrs.py)
- Rigid body docking: 1execute_haddock.sh
- Flexible docking: relax.sh
- RAbD design: rabd_design.sh
- Identify the light chain Fv gene: identify_lightchain.sh
- Binding energy calculation: run_ddg.sh (with ddg.xml)
- ESM embedding generation: run_esm.sh
- Training neural network: esm2.py / esm2_transfomer_lightning.py / esm2.ipynb

A REAMDE.md file in the code repository summarizes the provided scripts.

**Packages and dependencies.**

- GROMACS 2021
- Rosetta 3.14
- HADDOCK 2.4
- Pyrosetta 2022
- ANARCI
- AbNumber
- ESM
- PyTorch 2

The complete list of dependencies are available on the shared code repository.

#### 2. Datasets

The datasets are publicly available at Mendeley Data, containing the following:

- Raw data of figures in the manuscript: paper_data_plots
- $\alpha$S structures used for docking and residues on $\alpha$S as restraints for docking: synuclein_for_docking
- L3D1 structures: lag3_domain1
- *apo* Fv ensemble used for docking: fv_apo_ensemble.tar.xz
- Sequences with binding affinities used for model training: models/training_sequences.txt
- For binding affinities: training_ddg.pt
- Trained model with weights: models/weights
- Top predictions of the models (evaluation): models/eval_*

The following data is readily available upon request:

- Raw molecular dynamics trajectories (in .xtc format) of the *apo* Fv antibodies.
- Docked poses of the Fv-$\alpha$S complexes.

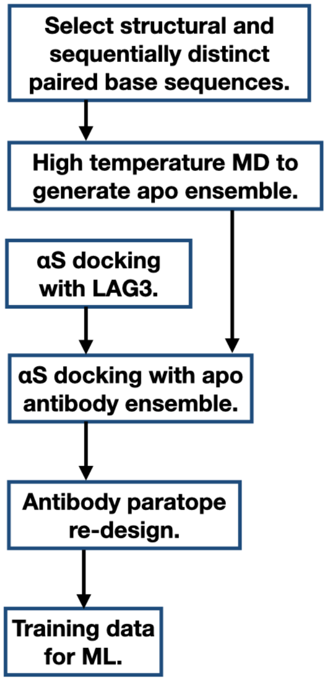


**Fig. S1.** A flowchart summarising the computational pipeline developed in this study.

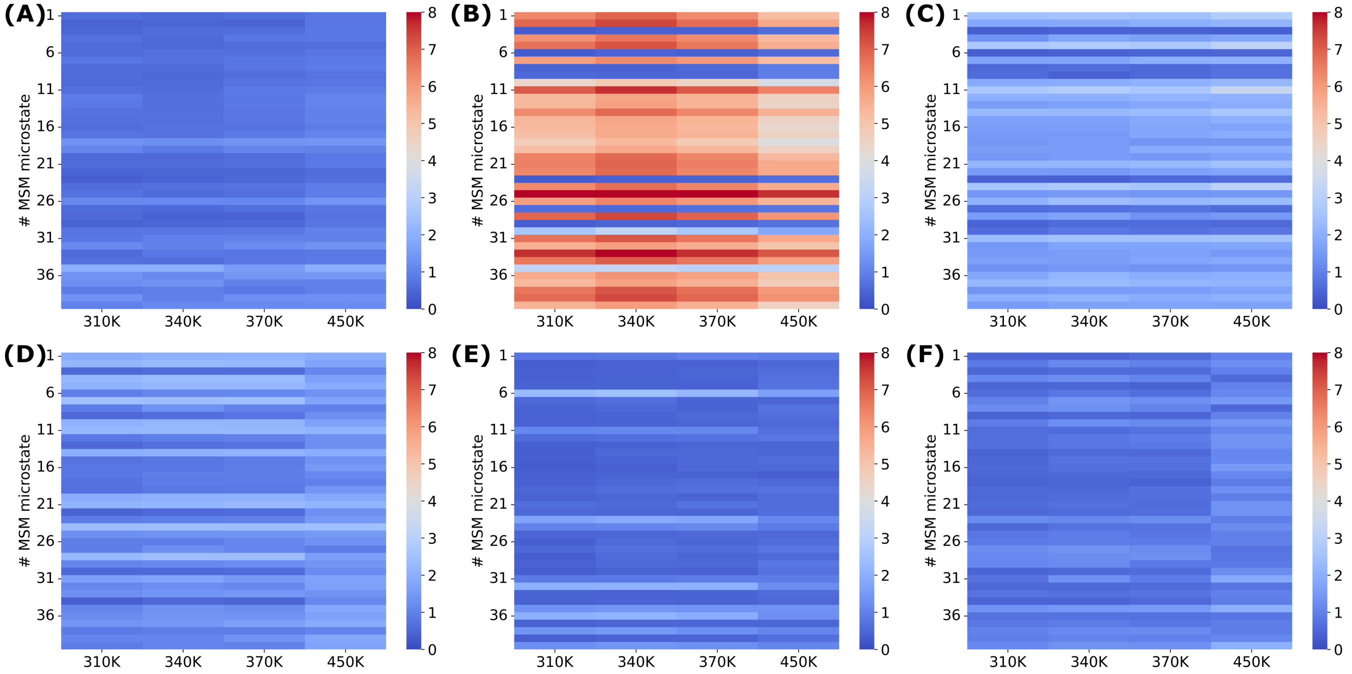


**Fig. S2. Comparison of CDR backbone RMSDs to microstates from a Markov-state model. A-F** The minimum CDR backbone RMSDs (in Å) of *apo* murine VEGF antibody, G6 (PDB ID: 2FJG) simulated at different temperatures is compared to the microstates dervied from a Markov-state model analysis. Subplots **A-C** depict the RMSDs corresponding to CDR H1-H3 and (**D-F**) correspond to CDR L1-L3 respectively. A $50\,\mathrm{ns}$ simulation was performed at each temperature starting from the *apo* state of the G6 Fv antibody.

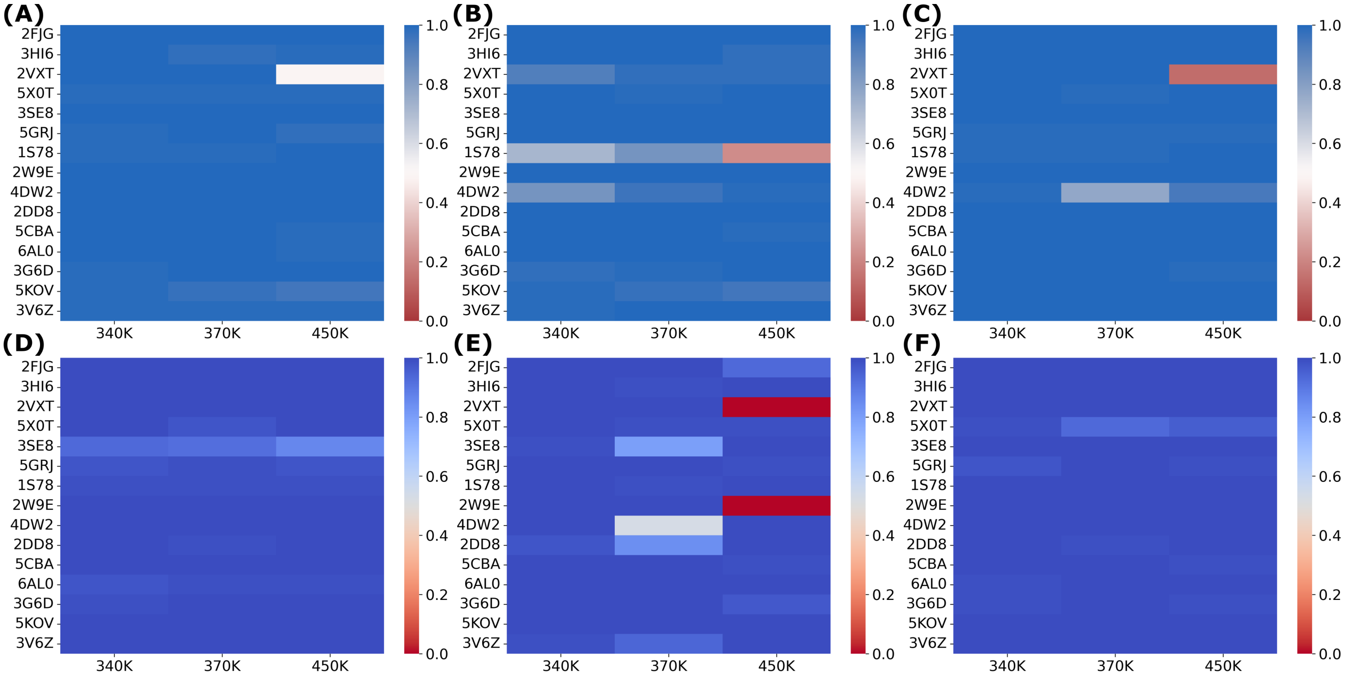


**Fig. S3. Similarily of trajectories generated at physiological temperature ($310\,\mathrm{K}$) to those generated at a higher temperature.** From the antibody set assembled by Guest et al. (1), the heatmaps highlight the fraction of MD trajectory frames generated at $310\,\mathrm{K}$ that are structurally similar to at least one frame from trajectories generated at higher temperatures ($340\,\mathrm{K}$, $370\,\mathrm{K}$, $450\,\mathrm{K}$). Two structures are similar if the combined CDR RMSDs are under $2\,\mathrm{\AA}$. MD trajectories of length $0.5\,\mu\mathrm{s}$ were generated at each temperature for comparison.

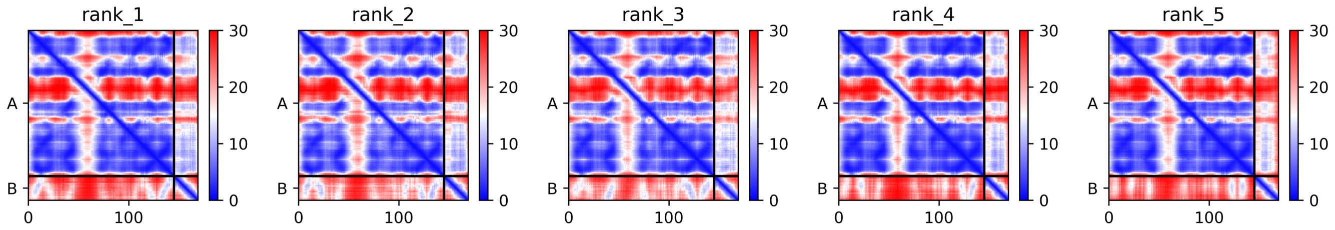


**Fig. S4. The predicted aligned errors (PAE) of the LAG3-$\alpha$S C-terminal peptide as obtained from the structure prediction from AlphaFold2 (2).**

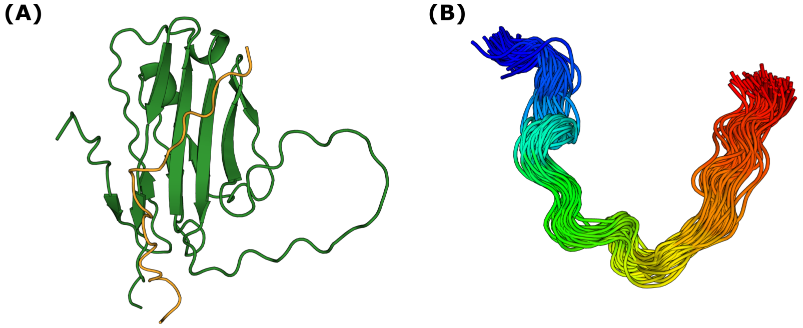


**Fig. S5. Visualization of structures used for docking. A** The complex of LAG3 Domain 1 (L3D1) with the C-terminus of $\alpha$S corresponding to the highest fraction of NMR contacts as determined by Zhang et al. (3). **B** An ensemble of the C-terminus of $\alpha$S, residues 118-140 (blue to red) generated via HADDOCK (4).

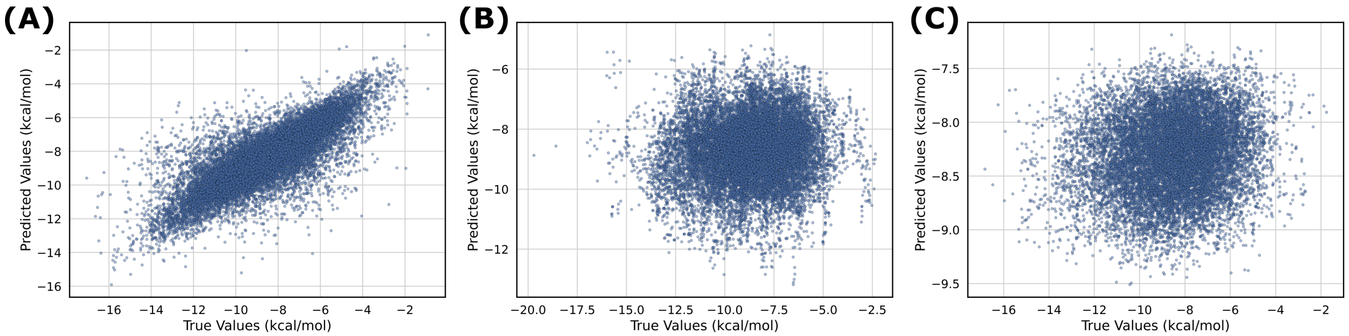


**Fig. S6. Model performance evaluation by measuring the correlation between the predicted and true binding affinity values. A** The correlation calculated using a feedforward network evaluated on a test set that shares > 80 % CDR sequence identity to the training set. **B** Correlation evaluation using a feedforward network on a test set that shares < 20 % CDR sequence identity to the training dataset. **C** Correlation plots using per-residue sequence embeddings generated from ESM2 ($150\ \mathrm{M}$ parameters) fed to a transformer layer on test dataset sharing < 20 % CDR sequence identity to the training dataset. The model architectures are depicted in Figure 5 (main text)

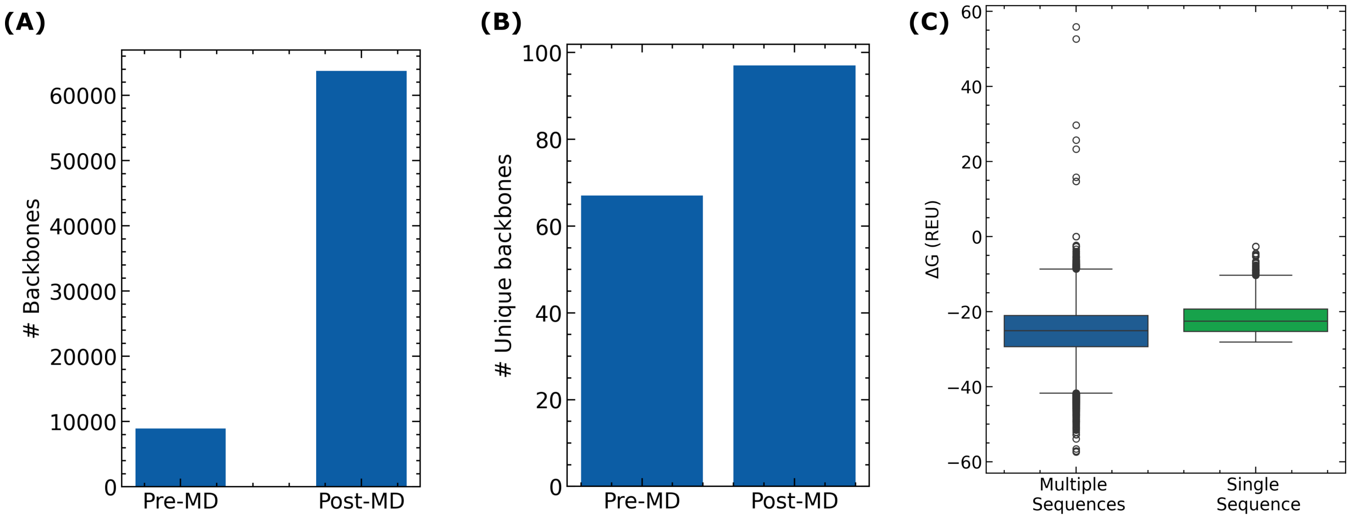


**Fig. S7. Enhancements in structural and sequence diversity. A** Post molecular dynamics, the number of backbones (after clustering) results in an average of 7 distinct conformations per Fv sequence. **B** Unique backbone folds as clustered by FoldSeek is plotted. MD yields larger number of unique folds. **C** The distribution of the Fv-$\alpha$S binding affinities ($\Delta$G) on taking a single Fv sequence (in green) compared to taking multiple sequences (in blue).
